# Structure of a zoonotic H5N1 hemagglutinin reveals a receptor-binding site occupied by an auto-glycan

**DOI:** 10.1101/2024.12.06.626699

**Authors:** Nicholas C. Morano, Yicheng Guo, Jordan E. Becker, Zhiteng Li, Jian Yu, David D. Ho, Lawrence Shapiro, Peter D. Kwong

## Abstract

Highly pathogenic avian influenza has spilled into many mammals, most notably U.S. dairy cows with several dozen human breakthrough infections. Zoonotic crossovers, with hemagglutinins mutated to enhance viral ability to use human α2-6-linked sialic acid receptors versus avian α2-3-linked ones, highlight the pandemic risk. To gain insight into these crossovers, we determined the cryo-EM structure of hemagglutinin from the zoonotic H5N1 A/Texas/37/2024 strain (clade 2.3.4.4b) in complex with a previously reported neutralizing antibody. Surprisingly, we found that the receptor-binding site of this H5N1 hemagglutinin was already occupied by an α2-3-linked sialic acid and that this glycan emanated from asparagine N169 of a neighboring protomer on hemagglutinin itself. This structure thus highlights recognition by influenza hemagglutinin of an “auto”-α2-3-linked sialic acid from N169, an *N*-linked glycan conserved in 95% of H5 strains, and adds “auto-glycan recognition” to the complexities surrounding H5N1 zoonosis.

**Highlights:**

- We report the structure of hemagglutinin from the zoonotic H5N1 strain A/Texas/37/2024 from clade 2.3.4.4b responsible for the current H5N1 outbreak
- Structural analysis revealed that each receptor-binding site of H5N1 A/Texas/37/2024 HA is bound to an “auto” sialic acid originating from glycan N169 on an adjacent protomer
- This structure provides new aspects of receptor binding for a highly pathogenic strain of avian influenza, and raises questions about the impact of auto-binding sialic acid, especially with respect to zoonosis

---

In recent years, a newly emerged highly pathogenic avian influenza virus strain of H5N1 (clade 2.3.4.4b) virus that was mostly confined to wild birds in Asia has spread globally and into new species of mammals, most notably dairy cows^1,2^. Since the onset of outbreaks in early 2024, 690 herds have been infected in 16 US states.^3^ Subsequent accounts of mammalian infection have also been reported, including in black bears^4^, foxes, skunks, raccoons, bobcats, and other terrestrial mammals^5^, as well as marine mammals such as the New England harbor and grey seal^6^. The spreading of this virus through wild birds and domestic poultry, to cows and other mammals raises the question of whether humans will be next. As of December 2024, 55 confirmed human breakthrough cases associated with clade 2.3.4.4b viruses spanning the United States and Canada have been reported. Thus far, zoonotic infections have been mild, though one individual – a Canadian teenager – was hospitalized in critical condition^7^. The historically high fatality rate of H5N1 infection in humans^8^, however, validates the threat associated with this current outbreak.

One factor that may underlie zoonosis is the preference of the H5N1 hemagglutinin for sialic acid receptors with either an α2-3 or an α2-6 glycosidic linkage to galactose. While avian influenzas tend to bind α2-3-linked sialic acids, human influenzas prefer α2-6 sialic acids, which are prevalent in the human upper respiratory tract^9^. Multiple reports have suggested, however, that clade 2.3.4.4b H5N1 hemagglutinin (HA) has little to no detectable affinity for α2-6, which could explain the lack thus far of human-to-human transmission^10,11^. Receptor specificity was recently probed using deep scanning mutagenesis experiments, which identify mutations at residues near the receptor binding site, which increase entry of H5N1 for cells expressing α2-6 glycans, and often simultaneously decrease entry into α2-3 expressing cells^12^. Many of the identified mutations are readily accessible by a single nucleotide mutation, suggesting H5N1 clade 2.3.4.4b may soon adapt to recognize α2-6 glycans. Further, mutations shown to enhance α2-6 cell entry^12^ have observed in break-through cases (**Figure S1**).

Although the source of the Canadian teenager’s case remains unknown, the HA of this virus is closely related to the Washington poultry farm outbreak and belongs to a new genotype, D1.1^13^, which is distinct from the cattle-related B3.13 genotype cases^13^ (**Figure S1**). Furthermore, the virus in the Canadian teenager’s case carries an A144T mutation, which has been shown to enhance α2-6 receptor cell entry^12^. Here, we used cryogenic electron microscopy (cryo-EM) to determine the structure of HA of H5N1 (A/Texas/37/2024) from clade 2.3.4.4b. To enhance resolution, we complexed this HA with 65C6, an antibody previously reported to neutralize H5N1^14^, which we show could also neutralize A/Texas/37/2024 at an IC50 of 0.067 ug/ml (**Figure S2A**). We used a soluble recombinant HA with an optimized Furin cleavage site and C-terminal appended T4 phage ‘Foldon’ domain^15^ (see methods). The Fab-HA complexed sample was frozen on grids, and cryo-EM data collected on a Titan Krios. A resolution of 2.68 Å was obtained (**Data S1, Table S1, Figure 1A**). We found antibody 65C6 to bind HA head as previously described^14^ (**Figure S2B-C)** and the recognized epitope to be conserved across H5N1 (**Figure S2D**).

**Figure 1.**
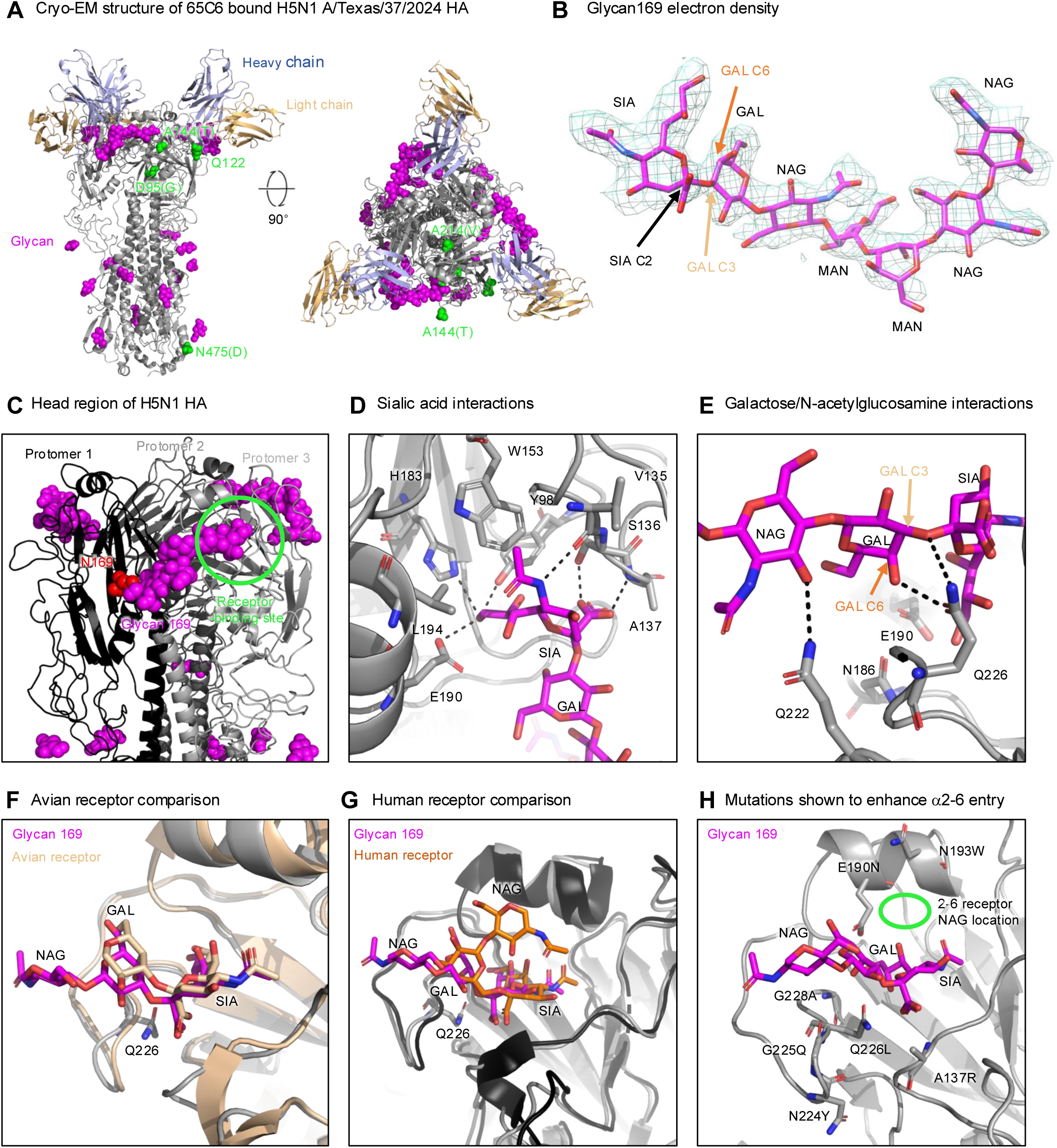
Cryo-EM structure of a zoonotic H5N1 HA in complex with 65C6 Fab reveals the receptor-binding site to be occupied by a sialic acid from N169. (A) Overall structure of HA-Fab complex is shown as a side view (left) and rotated to show a top view (right). Residues where mutations that enhance α2-6 cell entry and have been found in zoonotic infections are shown in green (note that Q122 is already mutated from leucine in A/Texas/37/2024 to an entry-enhancing glutamine). Glycans (magenta) are shown in all atom format, for each of the seven saccharides of N169 as well as for each of the four protein-proximal NAGs in the HA stem. (B) The cryo-EM reconstruction density is shown for the glycan chain emanating from N169 on each protomer. Glycan density was sufficiently resolved to unambiguously place each saccharide moiety in the N169 glycan. (C) Head region of HA. Each HA protomer colored a different shade of grey or black. Glycan169 stretches across the head region and terminates in the receptor-binding site of an adjacent protomer (green circle). (D) Interactions of the sialic acid of glycan N169 with the receptor-binding site. Hydrogen bonds are denoted by dotted black lines; residues that make substantial contact with sialic acid are labeled. (E) Interactions of galactose and N-acetylglucosamine. These glycans are coordinated by hydrogen bonds with glutamines at positions 222 and 226. (F) Overlay of auto glycan N169 and a homologous H5 HA structure (tan) bound to an avian receptor analogue (tan PDB:5Z88) demonstrates near identical positioning of sialic acid, and similar positioning of galactose, indicating overall binding mode between glycan N169 and an avian-receptor analogue is similar. (G) Overlay of auto glycan N169 and a homologous H2 HA structure (black) bound to a human receptor analogue (orange PDB:8TP2) indicates near identical positioning of sialic acid, with galactose in a similar region but at perpendicular angle due to altered glycosidic linkages; this results in the preceding N-acetylglucosamines being positioned differently. (H) Location of residues previously reported to enhance the ability of H5N1 to use the human α2-6-lined sialic acid as a receptor. Mutations labeled indicated the most drastic mutation reported by Dadonaite, et al. Most do not interact with sialic acid and are near the galactose residue, or on the strand which contains residue 226, for which mutations to alanine, asparagine, phenylalanine, valine and leucine enhance α2-6 usage. See also Figures S1 and S2, Data S1, and Table S1.

Surprisingly, the cryo-EM density indicated that the receptor-binding site was occupied by a ligand. The density for this ligand was at approximately 2.5-3.0 Å resolution (**Data S1 D-E**) allowing for unambiguous placement of seven saccharides NAG-NAG-MAN-MAN-NAG-GAL-SIA (**Figure 1B**), a seven membered chain which commonly occurs in the respiratory tissue of birds^11,16^. Density could only be fit with an α2-3 glycosidic linkage between the galactose and sialic acid (**Figure 1B**). The glycan chain originated from N169 and extended across the HA head, and sialic acid occupied the receptor-binding site on an adjacent protomer (**Figure 1C**).

Sialic acid contacted residues Y98, V135, S136, A137, W153, H183, E191, and L194, which collectively comprised the receptor-binding site (**Figure 1D**). The carbon-9 alcohol made three hydrogen bonds with Y98, H183, and E191, while the AcNH group bonded with the backbone carbonyl of V135. The carboxylic acid bonded with the sidechain of S136 and the nitrogen backbone of A137. Hydrophobic interactions were mediated largely by W153 and L190 (**Figure 1D**). The galactose made substantial surface area contacts with N182, E190, and Q226. The position of the galactose was coordinated by hydrogen bonding with Q226. The Q226 side chain bonded with an ether glycosidic linkage and the alcohol of carbon 6. The preceding N-acetyl-glucosamine hydrogen bonded with the side chain of Q222 (**Figure 1E**).

Comparison of the bound auto-glycan with the structure of a highly homologous H5 HA bound to an avian receptor analogue (PDB 5Z88) demonstrated general structural agreement of the sialic acid and galactose, indicating a shared binding mode between the auto-glycan, and receptors used for viral entry (**Figure 1F**). Comparison with a highly homologous H5 HA bound to a human receptor analogue (PDB 2WR4)^17^ demonstrated similar positioning of the sialic acid and galactose, but rotation of this galactose positioned the preceding N-acetylglucosamine at a ∼90° angle relative to the corresponding residue of the avian receptor. Dadonaite et. al identify multiple residues for which mutation can significantly enhance α2-6 cellular entry^12^. The most noteworthy of these mutations occurs at position Q226, which coordinates the galactose reside (**Figure 1H**).

Though the N169 glycosylation site was conserved in 95% of H5 HAs (**Figure 2A**), binding of this glycan to the receptor-binding site has rarely been observed. Reasons for the lack of glycan N169 auto-binding in structural biology experiments include (i) producing HA in insect cells or GNTi(-/-) cells that do not yield sialylated glycans^18–23^, (ii) HA is exposed to high concentrations of exogenous receptor, which may displace glycan 169^24^, or (iii) HA has mutations that inhibit sialic acid binding (e.g., Y98F)^25^ (**Figure 2B**). Indeed, crystal structures of HA produced in insect cells from H5N1 (A/Texas/37/2024) clade 2.3.4.4b were published after submission of this paper, and none of these showed an auto-bound glycan.^26^ Also, we note one structure published in October 2024 that does observe an auto-bound N169 glycan in an H5N8 strain^27^. Comparison of this structure with the structure of A/Texas/37/2024 HA showed general agreement in positioning of sialic acid, but variable locations for the rest of the saccharide chain (**Figure 2C**). Entropy analysis indicated that the surface below the extended glycan chain was highly variable, which may affect auto binding ability and positioning of the glycan chain (**Figure 2D**). Lastly, the auto-binding of N-linked glycan by H5N1 hemagglutinin may function as a glycan shield, as the “wrapping” from one protomer to another masks more protein surface (∼500 Å^2^) than a typical *N*-linked glycan. We calculated the glycan shielding of H5 by computational simulations of the masking glycan, and observed substantial masking of the receptor binding site, much more masking than would occur with a high mannose glycan (**Figure 2E**).

**Figure 2.**
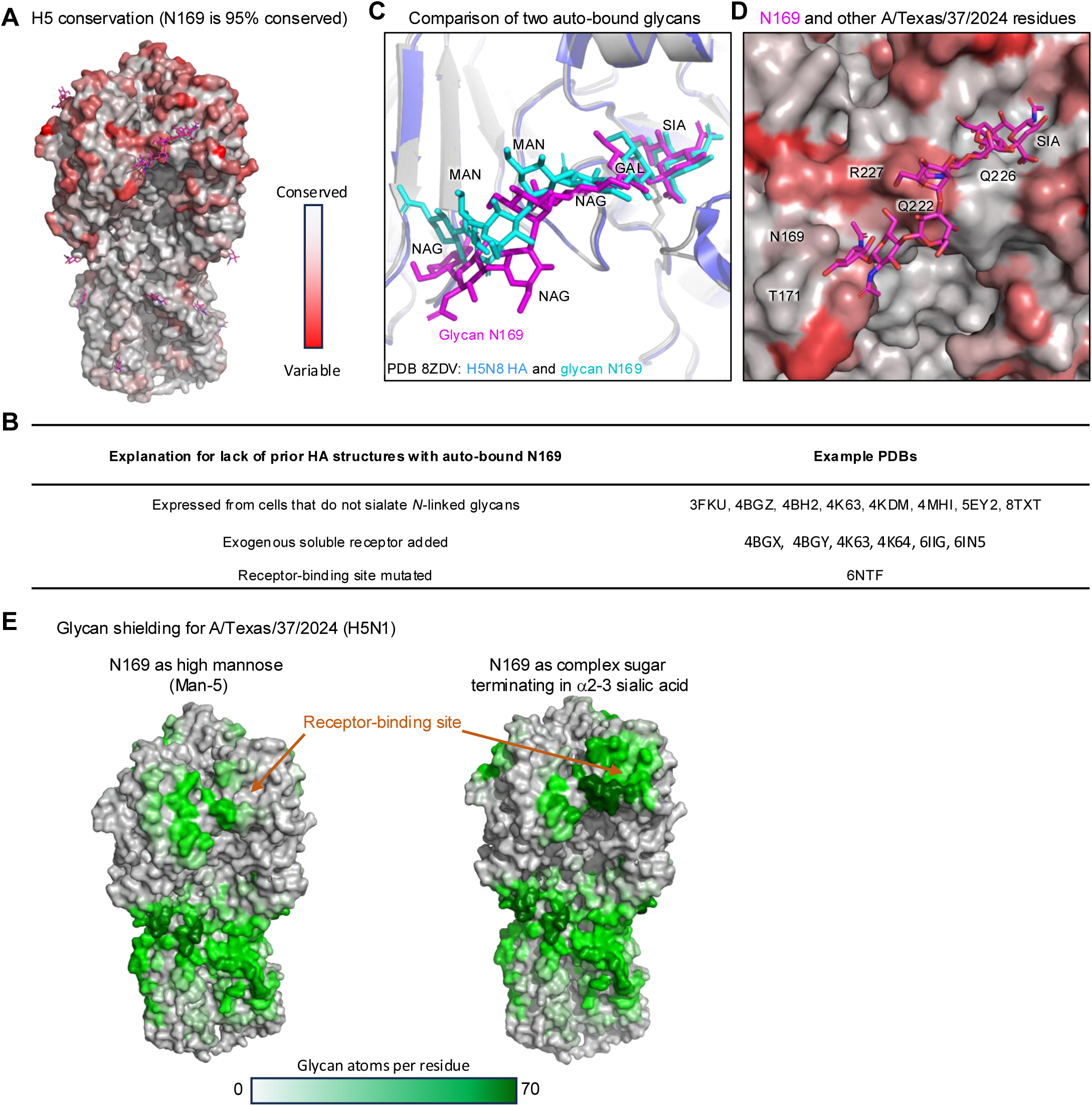
N169 conservation, explanation for lack of prior auto-binding, and impact on glycan shielding. (A) H5 conservation mapped to the surface of HA from A/Texas/37/2024 strain. Ordered saccharides are shown as magenta sticks. (B) Summary of representative H5 HA structures in the PDB. All observed to be N-linked glycosylated at residue 169. (C) Comparison of HA from H5N1 A/Texas/37/2024 (HA in grey, N169 in magenta) with HA from H5N8 (PDB 8ZDV) (HA in blue, N169 in cyan). (D) Close-up of the N169 glycan, with surface showing H5 conservation (with entropy scale the same as in panel A). The sialic acid-binding site, as well as N169, T171, and Q126, are highly conserved. The surface below the extended N169 glycan, however, is variable, likely influencing glycan positioning and degree of ordering. (E) Glycan shielding of H5 A/Texas/37/2024 strain as calculated with N169 as a high mannose glycan (Man5) (left) or as a α2-3 sialylated complex glycan (right). See also Figure S1, Data S1, and Table S1.

While we describe the structural, conservation, and glycan shielding aspects of the N169 auto-bound glycan, the functional impact of N169 on virion binding to cell-surface receptors or on virion egress from infected cells remains unclear. In terms of cell-surface receptor binding, a preloaded auto-bound glycan would need to be displaced to allow HA to bind host receptors on target cells. In terms of viral egress, an auto-bound influenza virion may have an advantage, as auto-binding of glycan N169 should competitively prevent binding to host-cell sialic acids, facilitating release from infected cells, a role normally performed by neuraminidase. An intriguing possibility is that auto-binding of glycan N169 does inhibit receptor attachment, until virion-associated neuraminidase is able to remove sialic acid from virion-associated HA. In this context, such competitive inhibition may act as a dispersal mechanism in which nearby cells are not infected as HA is auto-bound, but – as virion-associated neuraminidase releases sialic acid from N169 – this time-delayed release from auto-binding could in principle allow virions to recognize more distal target cells. It will be fascinating to delineate the precise physiological role(s) of glycan N169, to complement the structural, conservation, and glycan shielding aspects of this auto-bound glycan in the structure of the zoonotic H5N1 HA from clade 2.3.4.4b.

## Acknowledgements

We thank S. Chong for assistance with neutralization and members of ADARC for scientific discussions and comments. Cryo-EM data collection was performed at the Columbia University Cryo-Electron Microscopy Center. This work was supported in part by Bill and Melinda Gates Foundation grant INV-016167 to L.S.

## Author contributions

N.C.M., Y.G., D.D.H., L.S., and P.D.K. conceived the study. N.C.M determined the cryo-EM structures and conducted structural analysis. Y.G. conducted phylogenetic analysis, entropy analysis, and antibody analysis. J.E.B. prepared cryo-EM grids and assisted in protein production and purification. Z.L. and J.Y. conducted pseudoviral neutralization experiments. D.D.H. supervised neutralization experiments. P.D.K. and L.S. supervised structural experiments. N.C.M., Y.G., L.S. and P.D.K. prepared the first draft of the manuscript, which all authors reviewed and edited. All authors agreed to submit the manuscript, read and approved the final draft, and assume full responsibility for its content, including the accuracy of the data and the fidelity of the analyses.

## Declaration of Interests

The authors declare no competing interest.

## STAR★Methods

### Resource Availability

#### Lead Contact

Further information should be directed to and will be fulfilled by Peter D. Kwong.

### Materials Availability

Requests for resources and reagents should be directed to and will be fulfilled by Peter D. Kwong. All new reagents are available by MTA for non-commercial research.

### Data and Code Availability

- Cryo-EM structures were deposited under PDB 9EKF. Associated cryo-EM map was deposited to EMDB under EMD-48120.
- This study did not generate new code.
- Any additional information required to reanalyze the data reported in this paper is available from the lead contact upon request.

### Phylogenetic Analysis

The HA amino acid sequences of H5N1 clade 2.3.4.4b were obtained from the GISAID database^28^. These sequences were aligned using MUSCLE^29^, and a maximum likelihood tree was built using MEGA11^30^, with the general time reversible (GTR + G) substitution model, and the A/American wigeon/South Carolina/AH0195145/2021 was used to root the phylogenic tree.

### Cell lines

Expi293F cells were from ThermoFisher Scientific Inc (cat# A14527). TZM-bl cells were from NIH AIDS Reagent Program (www.aidsreagent.org, cat# 8129). The above cell lines were used directly from the commercial sources and cultured following manufacturer suggestions as described in Method Details below.

### Pseudoviral Neutralization Assay

Pseudotyped H5N1 (pseudoviruses) were packaged with NL4-3-LucΔenv backbone as previously described^31,32^. Briefly, the plasmids, hemagglutinin (HA), neuraminidase (NA), and NL4-3-LucΔenv backbone were co-transfected into HEK293T cells with lipofectamine 3000 following the manufacturer instructions. 8 hours post-transfection, the supernatant was replaced with fresh medium. After 48 hours, the supernatant was collected by centrifugation, aliquoted and store at −80°C. Prior to use in neutralization assays, the pseudoviruses were titrated. For the neutralization assay, serial diluted antibodies were prepared in 96-well plates, starting at a concentration of 10 μg/mL. Subsequently, titrated pseudoviruses were added and incubated at 37°C for 1 hour. Following this, 5,000 MDCK cells were seeded into each well and incubated for another 48 hours. Cells were lysed and luminescence signals were measured using a luciferase assay kit (Promega) on a SoftMax Pro v7.0.2 (Molecular Devices). The IC50 values were calculated by fitting a five-parameter non-linear dose-response curve using GraphPad Prism 9.2.

### 65C6 Antibody Production

For antibody expression, heavy chain and light chain variable regions of antibody genes were synthesized (Genscript) and subcloned into corresponding gWiz vectors. The heavy chain was cloned into an antibody expression vector with an HRVC3 protease site in the hinge region for FAB production. For antibody production, variable regions of the isolated antibody heavy chain genes were subcloned into corresponding pVRC8400 vectors, in which a HRV3C cleavage site was inserted in the hinge region. The heavy and light chain plasmids were mixed at a ratio of 1:1 and transfected into Expi293F cells using Expifectamine 293 transfection kit. At day 5 post transfection, the culture supernatant were harvested, clarified, and loaded onto a PrismA columns (Cytivia). The columns were washed with PBS and the antibodies were eluted with Ig Elution Buffer (Peirce), which was immediately neutralized with 1 M Tris, pH 8.0. Antibody was then purified by size exclusion chromatography using an S200 Increase 16/60GL Column (Cytiva) into 1X PBS, pH 7.5.

### 65C6 Fab Production

65C6 Fab was produced using HRV 3C protease kite (Pierce), Fc and uncut Ig were removed using protein A resin (Cytiva) and Fab was further purified by size exclusion chromatography using an S200 Increase 16/60GL Column (Cytiva).

### HA Construct Design and Protein Production

The sequence of H5N1 HA (A/Texas/37/2024) was engineered to replace the Furin cleavage site (REKRRKR) with (RRRRRR) for enhanced Furin cleavage. The transmembrane and cytosolic regions were removed and replaced with a 2X GSS linker, followed by a T4 Fold domain (GYIPEAPRDGQAYVRKDGEWVLLSTFL), glycine residue, 10x HIS tag, GGSG linker, and then AVI tag: MENIVLLLAIVSLVKSDQICIGYHANNSTEQVDTIMEKNVTVTHAQDILEKTHNGKLCDLNGVKPLILKDCSVAGWLLGNPMCDEFIRVPEWSYIVERANPANDLCYPGSLNDYEELKHMLSRINHFEKIQIIPKSSWPNHETSLGVSAACPYQGAPSFFRNVVWLIKKNDAYPTIKISYNNTNREDLLILWGIHHSNNAEEQTNLYKNPITYISVGTSTLNQRLAPKIATRSQVNGQRGRMDFFWTILKPDDAIHFESNGNFIAPEYAYKIVKKGDSTIMKSGVEYGHCNTKCQTPVGAINSSMPFHNIHPLTIGECPKYVKSNKLVLATGLRNSPLRRRRRRGLFGAIAGFIEGGWQGMVDGWYGYHHSNEQGSGYAADKESTQKAIDGVTNKVNSIIDKMNTQFEAVGNNLERRIENLNKKMEDGFLDVWTYNAELLVLMENERTLDFHDSNVKNLYDKVRLQLRDNAKE LGNGGCFEFYHKCDNECMESVRNGTYDYPQYSEEARLKRESSGSSGYIPEAPRDGQAYVRKDGEWVLLSTFLGHHHHHHHHHGGSGLNDIFEAQKIEWHE

Codon optimized DNA was synthesized by Genscript, and subcloned into the gWiz vector backbone for mammalian transient transfected. Soluble, fully cleaved H5 HA trimers were produced by transient co-transfection with Furin enzyme in Expi293F cells (Life Technologies) using Expi293 Transfection reagent. After 5 days at 37 °C, culture supernatants were harvested by centrifugation and concentrated 5-fold by Tangential Flow Filtration. The recombinant HA trimer was captured by Ni-NTA (Sigma-Aldrich) through a C-terminal 6xHis-tag. The imidazole eluant was combined 1:1 (v/v) with saturated ammonium sulfate, centrifuged at 4 °C, and pellet removed. The supernatant was dialyzed against 500 mM NaCl, 50 mM Tris pH 8, and purified by size exclusion chromatography on a Superdex 200 Increase 10/300 GL column (Cytiva).

### Cryo-EM Grid Preparation

Samples for cryo-EM grid preparation were produced by first mixing 15 µl of purified H5H1 HA (A/Texas/37/2024) at .5 mg/ml with 4 µl of 65C6 Fab at 3 mg/ml. Complex was adjusted to have a final concentration of 0.005% (w/v) n-Dodecyl β-D-maltoside (DDM) to prevent preferred orientation and aggregation during vitrification and incubated on ice for 20 minutes. Cryo-EM grids were prepared by applying 3 μL of sample to a freshly glow discharged carbon-coated copper grid (CF 1.2/1.3 300 mesh). The sample was vitrified in liquid ethane using a Vitrobot Mark IV with a wait time of 30 s, a blot time of 3 s, and a blot force of −5.

### Image Processing

Cryo-EM data were collected on a Titan Krios operating at 300 keV, equipped with a K3 detector (Gatan) operating in counting mode. Data were acquired using Leginon^33^. The dose was fractionated over 50 raw frames. For all structures, the movie frames were aligned and dose-weighted using cryoSPARC 3.4^34^; the CTF estimation, particle picking, 2D classifications, ab initio model generation, heterogeneous refinements, homogeneous 3D refinements and non-uniform refinement calculations were carried out using cryoSPARC 3.4.

### Atomic Model Building and Refinement

For structural determination, a starting model of the antibody Fab 65C6 was obtained from the previously deposited crystal structure of 65C6 (PDB 5DUM). For HA, the crystal structure of a computationally optimized H5 influenza hemagglutinin (PDB 6NTF) was used. The Fab and HA starting models where docked into the cryo-EM density map using UCSF Chimera to build an initial model of the complex. The model was then manually rebuilt to the best fit into the density using Coot and refined using Phenix^35^. Interface calculations were performed using PISA. Structures were analyzed and figures were generated using PyMOL (http://www.pymol.org) and UCSF Chimera). Final model statistics are summarized in Data S1 and Supplementary Figure S1.

### HA Protein Sequence Conservation Analysis

The HA protein sequences for conservation analysis were downloaded from GISAID, including all available H5 genomes with an HA sequence deposited. Low-quality sequences were removed, and duplicates were filtered using USEARCH^36^. The sequences were aligned using MUSCLE with default parameters. The conservation of each RBD residue was calculated using the entropy function of the bio3d R package (H.norm column) based on the HA sequence alignment. Sequence entropy was visualized on the HA structure using PyMOL version 2.5.4.

### Glycan Density Analysis

Man5 glycans were added using GlycoSHIELD^37,38^ to generate 10 conformations for each glycan. The N169 glycan with sialic acid were the replace the Man5 on N169 for each conformation of fully glycosylated structures. Subsequently, GLYCO^39^ was employed to calculate the glycan density per residue on the trimer structures and visualized using PyMOL version 2.5.4.

## Supplemental Files

**Figure S1.**
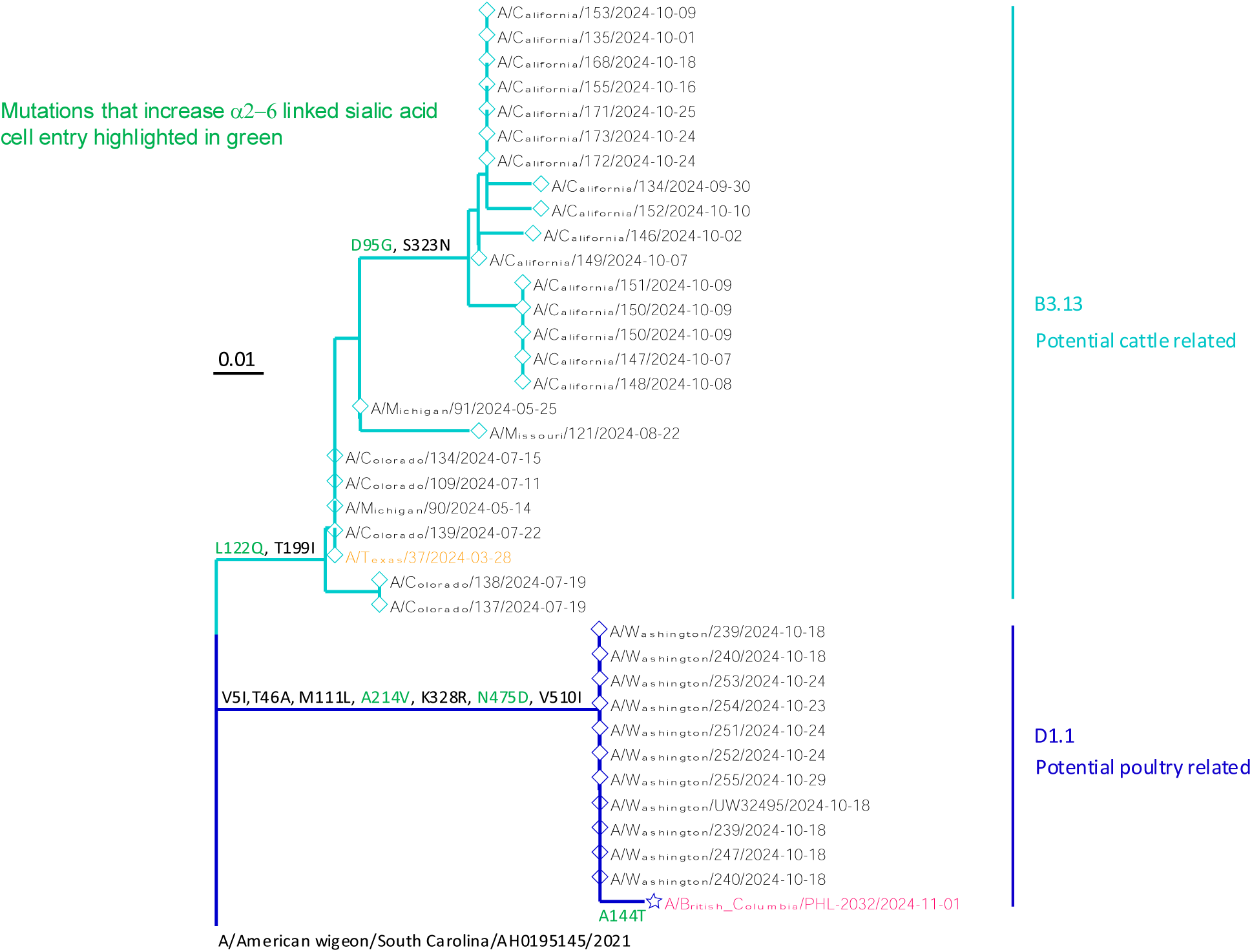
Phylogenetic tree of zoonotic clade 2.3.4.4b H5N1 infections and mutations that enhance α2-6 cell entry, related to Figures 1 and 2. Viral sequences isolated from a zoonotically infected Canadian teenager and the first case in the Americas in 2024 are highlighted in magenta and orange, respectively. Position numbering for mutations is based on mature H3 numbering.

**Figure S2.**
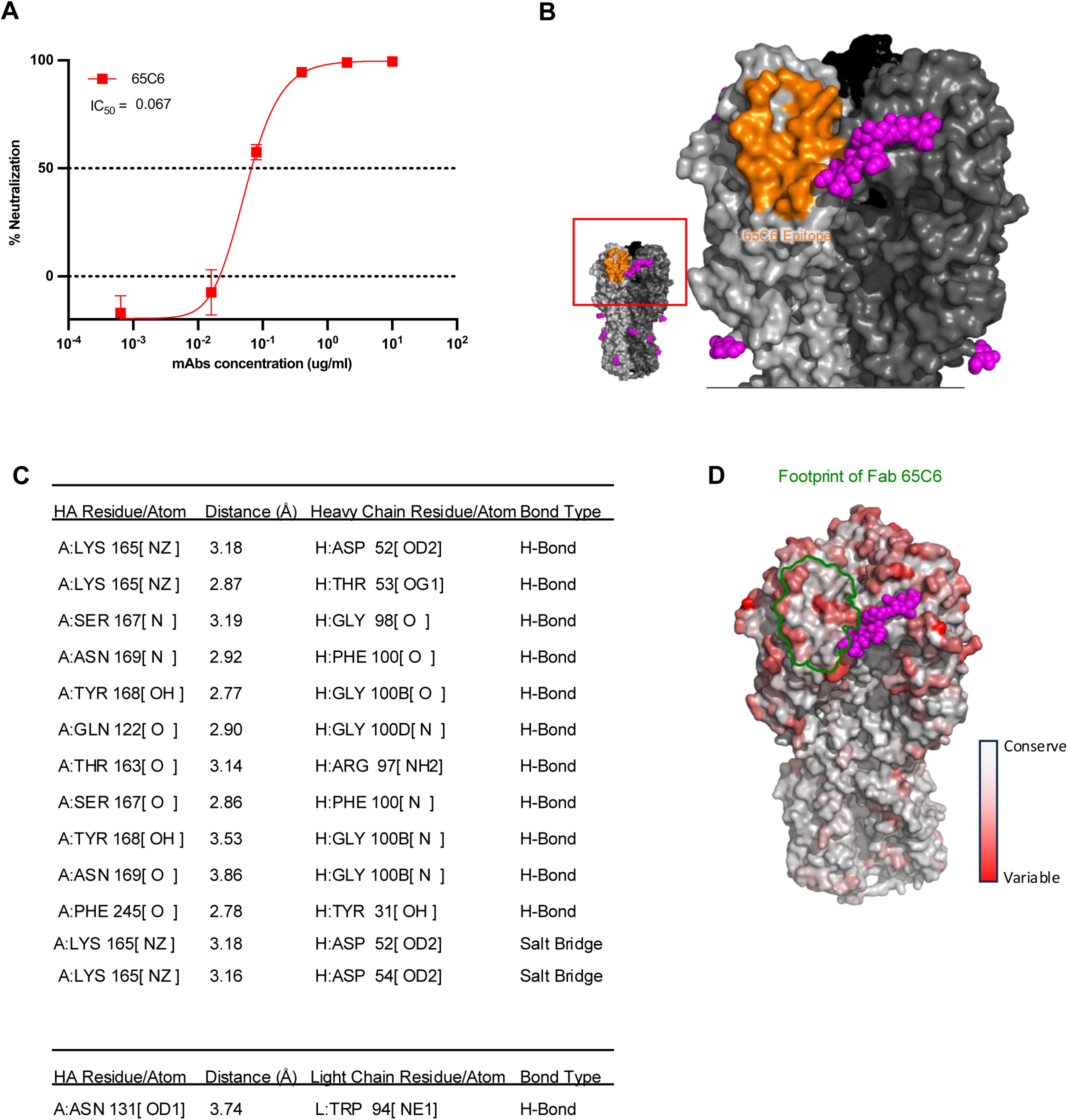
65C6 neutralization and cryo-EM details of the H5N1 HA in complex with 65C6 Fab, related to Figure 1. (A) Neutralization of A/Texas/37/2024 pseudo type virus by monoclonal antibody 65C6. (B) Epitope footprint of Fab 65C6 shown on one protomer of HA in orange, glycan 169 shown in magenta. (C) Pisa analysis of H bonds and salt bridges formed at 65C6 epitope and paratope (D) Footprint of Fab 65C6 shown on one protomer of HA outlined in green. Surface of HA indicates surface entropy, showing that 65C6 binds to a conserved site on the HA head. Glycan 169 shown in magenta.

**Table S1.** Cryo-EM data collection, refinement, and validation statistics, related to Figures 1 and 2.

|  |  |
| --- | --- |
|  | A/Texas/37/2024<br>H5N1 HA<br>(EMD-48120)<br>(PDB 9EKF) |
| **Data collection and processing** |  |
| Magnification | 105000 |
| Voltage (kV) | 300 |
| Electron exposure (e-/Å<sup>2</sup>) | 58 |
| Defocus range (µm) | .8-2 |
| Pixel size (Å) | 0.825 |
| Symmetry imposed | C3 |
| Initial particle images (no.) | 315964 |
| Final particle images (no.) | 206541 |
| Map resolution (Å) | 2.99 |
| FSC threshold | .143 |
| **Refinement** |  |
| Initial model used (PDB code) | 6NTF |
| Model resolution (Å) | 2.68 |
| FSC threshold | .143 |
| Model composition |  |
| Non-hydrogen atoms | 17531 |
| Protein residues | 2145 |
| Glycan residues | 33 |
| B factors (Å<sup>2</sup>) |  |
| Protein | 79.69 |
| Glycans | 49.10 |
| R.m.s. deviations |  |
| Bond lengths (Å) | 0.006 |
| Bond angles (°) | 1.139 |
| Validation |  |
| MolProbity score | 1.76 |
| Clashscore | 4.34 |
| Poor rotamers (%) | 2.15 |
| Ramachandran plot |  |
| Favored (%) | 95.80 |
| Allowed (%) | 4.20 |
| Disallowed (%) | 0.0 |

**Data S1.**
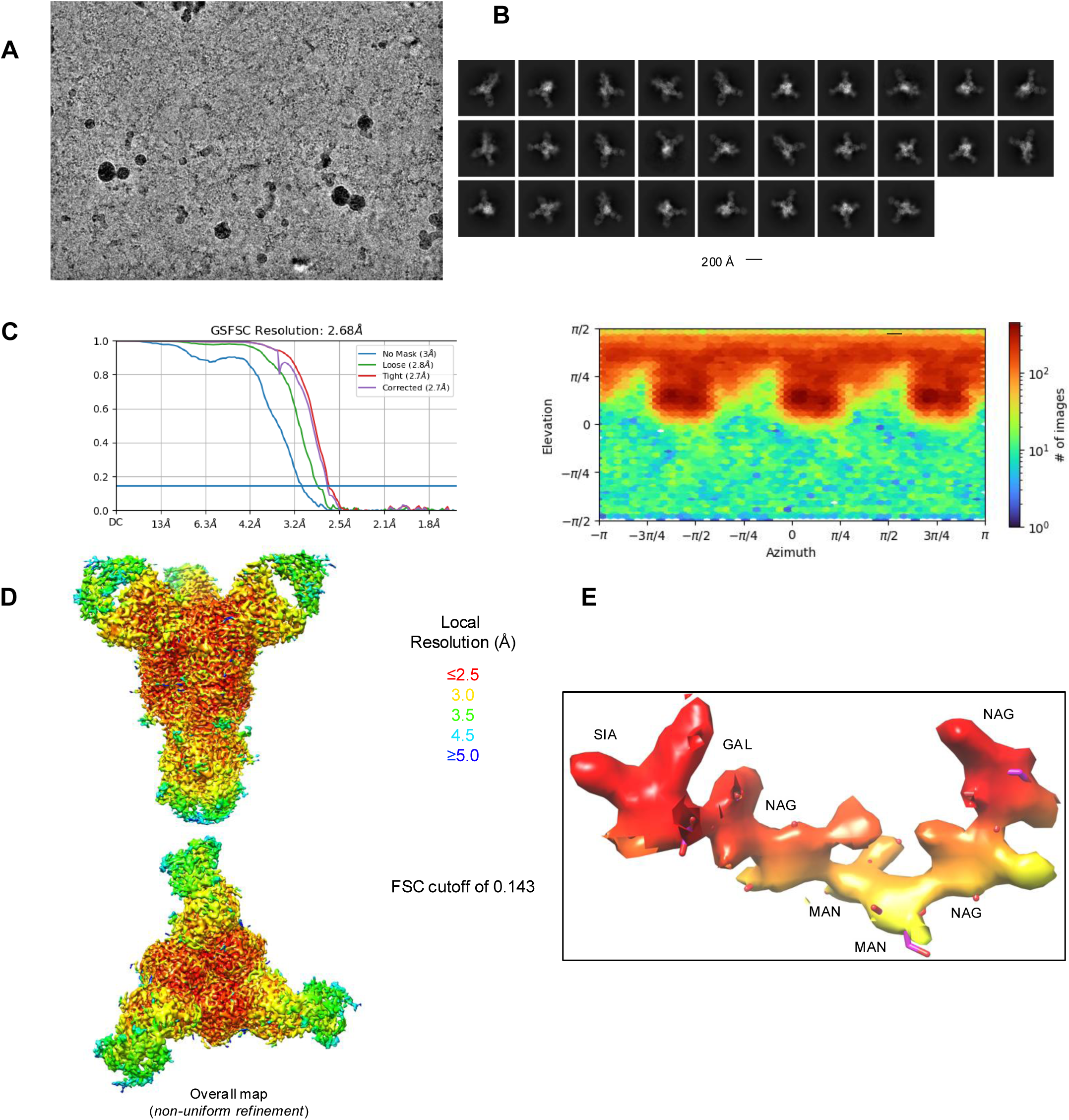
Cryo-EM details of the H5N1 HA in complex with 65C6 Fab, related to Figures 1 and 2. (A) Representative micrograph. (B) Representative 2D class averages are shown. (C) The gold-standard Fourier shell correlation resulted in a resolution of 2.68 Å for the overall map using non-uniform refinement with C3 symmetry (left panel); the orientations of all particles used in the final refinement are shown as a heatmap (right panel). (D) The local resolution of the final overall map is shown contoured at 0.202. Resolution estimation was generated through cryoSPARC using an FSC cutoff of 0.143. (E) Representative density with resolution coloring is shown for Glycan169.

## Notes

### Competing Interest Statement

The authors have declared no competing interest.

